# Towards Personalized Epigenomics: Learning Shared Chromatin Landscapes and Joint De-Noising of Histone Modification Assays

**DOI:** 10.1101/2025.03.04.641154

**Authors:** Tanmayee Narendra, Giovanni Visonà, Crhistian de Jesus Cardona, James Abbott, Gabriele Schweikert

**Author notes:** Contributing authors.

## Abstract

Epigenetic mechanisms enable cellular differentiation and the maintenance of distinct cell-types. They enable rapid responses to external signals through changes in gene regulation and their registration over longer time spans. Consequently, the chromatin landscape, which is the overall organization and biochemical state of chromatin, exhibits both cell-type and individual specificity and contributes to phenotypic diversity. Genomic distributions of chromatin features are typically measured using ChIP-Seq and related methods. However, these measurements are subject to substantial biases introduced by the chromatin landscape itself. Here, we introduce DecoDen, which uses measurements of several different histone modifications, to simultaneously learn shared chromatin landscapes while de-biasing individual measurement tracks. We demonstrate DecoDen’s effectiveness on an integrative analysis of histone modification patterns across multiple tissues in personal epigenomes.

## Introduction

Although the cells of a multicellular organism contains nearly identical copies of DNA, they vary widely in form, shape, and function. One of the many ways cells regulate function is by creating distinct chromatin environments that enable the precise execution of cell-type-specific tasks: A complex array of epigenetic factors dynamically modify the physical properties of DNA, introducing chemical changes to both the DNA molecule itself and to histone proteins, around which DNA is wound. Through these modifications, epigenetic mechanisms either facilitate or restrict access to specific genomic regions, influencing interactions between transcription factors and their corresponding DNA substrates - ultimately enabling the wide range of cellular phenotypes observed across different cell types.

Epigenetic changes play vital roles throughout all stages of life: During development, proliferating cells encode essential information in their chromatin states [1], which is passed down to daughter cells. Cellular differentiation is accompanied by tightly regulated adjustments to the chromatin landscape [2]. Throughout an organism’s lifetime, as cells transition into quiescent states, epigenetic changes mediate the integration of environmental signals, molecular cues, and cellular metabolism to maintain homeostasis and preserve cell identity [3]. Eventually, chromatin modifications are involved in regulating aging processes [4], and aberrant chromatin reprogramming is an early hallmark of tumor development [5].

Obtaining cellular snapshots of epigenomic landscapes remains expensive and time-consuming, requiring extensive experimental efforts, with large numbers of individual assays to record histone modifications individually. Over the last decade international efforts have been undertaken to create tissue and cell-type specific epigenomic reference maps, for example, the Roadmap Consortium established epigenomic references for 111 tissue and primary cell types, focusing on a core subset of epigenomic marks [6]. The primary experimental technique used is chromatin immunoprecipitation sequencing (ChIP-Seq) [7–9]. Despite being widely adopted, ChIP-Seq has several limitations. The chromatin fragments used for ChIP-Seq experiments are generated by sonication, which has an inherent bias towards accessible chromatin [10]. Further, ChIP-Seq uses formaldehyde cross-linking which results in biased recovery of accessible chromatin [11]. Together, this may have an unequal impact on different histone marks (see Figure S4 in Supplementary Material.) [12, 13], and false positive phantom peaks can be detected in active promoter regions [14].

While reference epigenomes have proven invaluable, they are insufficient for identifying functional important or disease-causing epigenomic variations. Individual-specific epigenomes are now emerging, with the EN-TEx resource providing post mortem data from 15 assays from four individuals across 30 tissues [15]. Additionally, computational methods have been developed to fill remaining experimental gaps through cross-cell type imputation of epigenomic profiles. We have earlier contributed to these efforts by developing a transformer-based method, eDICE [16], which was positively assessed on reference epigenomes in the ENCODE Imputation Challenge [17]. Using eDICE, we have also shown that it is possible to train on data from one individual and use transfer learning to predict histone modification patterns in a target individual, where only limited measurements are available. This proof of concept illustrates the potential for imputation of tissue specific histone modifications from readily accessible samples.

However, despite these technical advancements and an overall acceleration in data production, the analysis and interpretation of the resulting data sets remains challenging. Notably, individual histone modifications are far from independent of each other, but exhibit non-linear, locally changing combinatorial arrangements. This is not surprising, as epigenetic factors simultaneously interpret and effect upon associated modification patterns. Indeed, the understanding of the dynamic interactions between individual epigenetic components is of particular importance. However, the resulting co-created chromatin landscape strongly impacts the local sensitivity and specificity of the measurements, thus introducing confounding shared biases. To understand the concerted cell-specific interaction between epigenetic factors, as well as to detect subtle changes in individual histone modifications between individuals, it is essential to efficiently and robustly separate the joint effect of the chromatin environment from the interrogated histone modifications itself. To this end, input control experiments are typically conducted that estimate the local effect of the chromatin environment. Typically, these are used computationally to correct for each assay track individually. However, for an effective separation of chromatin bias from the local distribution of histone modification enrichment, the bias has to be correctly estimated genome-wide. This is rarely the case. Typically, only 40-50 million fragments are sequenced in the control experiment, such that large proportions of the human genome remain not represented or undersampled, with sometimes 16% of the genome not covered at all (see Figures S1, S2 and S3 in Supplementary Material for examples). As a consequence, input control samples alone are not sufficient to correct the measurements and may even introduce additional noise. Without correct debiasing of the measured data, experimental results can be misinterpreted and computational imputation methods which utilize existing data to perform in-silico predictions of missing histone modifications will invariably learn the data inherent biases, and propagate them into the computational predictions.

Despite these challenges, the potential for personalized epigenomics, in which individual-specific chromatin differences are identified in an unbiased manner, presents a compelling case for improving the precision of histone modification profiling and imputation. A deeper understanding of inter-individual chromatin differences is essential for predicting disease progression and developing targeted therapies, underscoring the need for more refined and precise methodologies in chromatin research.

We introduce DecoDen (Deconvolve and Denoise), a new method to remove cell-type-specific bias for samples of different histone modifications derived from the same cell or tissue samples. This method leverages different assays and jointly analyses data to learn a shared chromatin bias common to all assays performed on a given sample. With simulated data we show that DecoDen accurately reconstructs hidden signals and demonstrate its superior performance through quantitative comparison with other methods. Applying DecoDen to a number of different experimental data sets, we demonstrate its unique ability to minimize confounding effects. Lastly, we test DecoDen by reanalysing multi-tissue personal epigenomes from the EN-Tex resource [18] to identify individual-specific differences in histone modification patterns. However, we find severe limitations in the experimental data quality and discuss improved experimental guidelines. DecoDen is available as an open-source tool at https://github.com/ntanmayee/DecoDen.

## Material & Methods

### The DecoDen Model

DecoDen is an integrative approach that shares information between multiple assays and samples from a given cell-type or tissue, to reconstruct hidden signals of different histone modifications in the context of a shared chromatin landscape. The DecoDen model is founded on three specific modelling assumptions, which are addressed in two consecutive steps of deconvolution and denoising.

1. The first assumption is that the observed treatment measurement consists of a target-specific signal and background reads, which originate from *independent* processes. In other words, the final observed measurement is the sum of a target-specific component and a background component.
2. Second, we assume that the process from which background reads are generated is shared across all assays. Further, this process is independent of their specific target, as long as they are performed on the same biological sample. This allows us to share information across multiple assays in order to improve upon a locus-specific background control estimate for the whole genome. The additive bias is addressed using a constrained form of Non-negative Matrix Factorization (NMF), which deconvolves all observed measurements into target-specific signals and a shared background. This step also identifies the coefficients for their linear combinations. These mixing parameters are related to the global specificity of the used antibodies and the observed signal-to-noise ratios of the individual measurements.
3. The third assumption is that the learned unspecific background purely captures biases such as local chromatin accessibility and sequencing mappability, which equally affect the target specific signals by locally amplifying or reducing the number of sequenced reads. We thus use the unspecific background to further remove confounding biases from the target-specific signals by applying Half-Sibling Regression (HSR) [19], an approach from causal inference that aims to reconstruct a latent quantity of interest.

A conceptual overview can be seen in Figure 1.

**Fig. 1:**
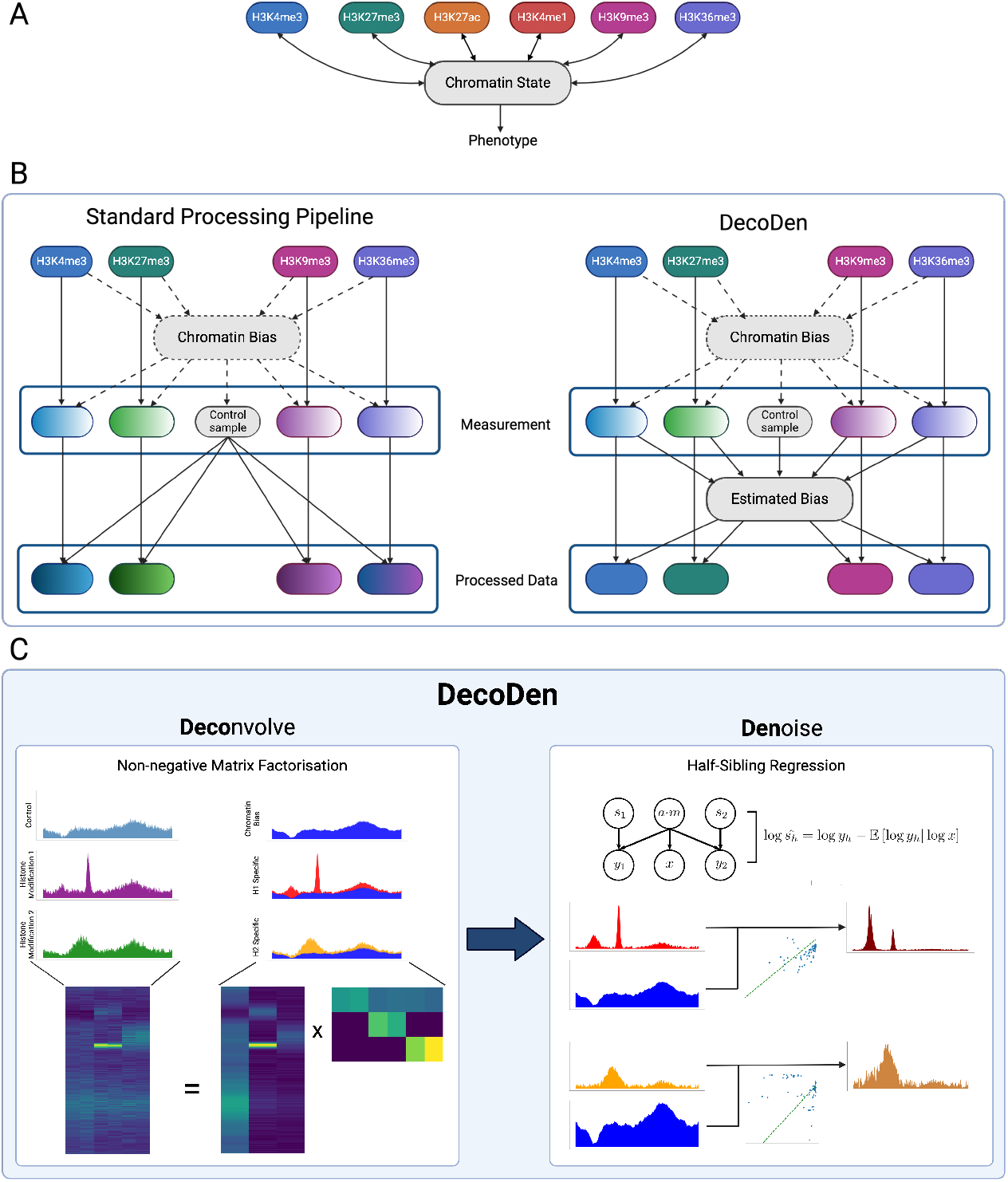
Schematic of DecoDen: (A) Histone modifications interact with each other via chromatin to jointly establish ‘chromatin states’ that determines the DNA’s accessibility to regulatory proteins, thereby influencing transcription and the observed phenotype. Measurement of histone modifications are confounded by this chromatin state. (B) In the standard data processing pipeline, the data for each modification is independently analysed and estimates are implicitly biased by the chromatin state (chromatin bias). With DecoDen, all samples are jointly analysed to estimate a chromatin bias which is consequently minimised from measurement samples. (C) In the first step of DecoDen, Non-negative Matrix Factorization (NMF) is used to separate the cell-type specific chromatin signal from the assay-specific signal of the di!erent histone modification measurements. Next, Half-Sibling Regression (HSR) is used to extract the true (hidden) enrichment pattern by minimizing the bias introduced by accessibility and mappability.

We can define the previous qualitative description with a suitable formalism (vectors are represented in bold typeface): Let us assume that local biases which depend on the DNA sequence alone, can be described with a vector **m** ∈ ℝ^*G*^, where *G* is the length of the binned genome. Cell-type-specific biases including chromatin accessibility are represented with **a** ∈ ℝ^*G*^ and **n** ∈ ℝ^*G*^ represents the nonspecific signal that captures any other process not included in the aforementioned biases.

The observed *r*-th replicate of a control (input) measurement is given by **c**^(*r*)^ ∈ ℝ^*G*^. The value measured at each genomic bin *b* can then be written as:

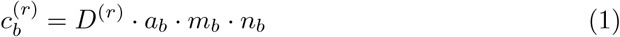

where *D*^(*r*)^ represents the replicate-specific scaling factor accounting for variable sequencing depth.

Further, we can describe the observed data 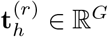 for an an experimental assay for the *h*-th histone modification and *r*-th replicate as:

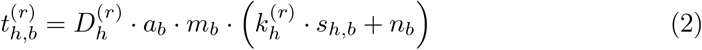

where again 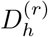 represents the scaling factor for the *r*-th replicate for modification *h. s*_*h,b*_ is the true (latent) signal enrichment for the *h*-th histone modification in genomic bin *b*, and 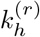 is a mixing parameter that captures the antibody specificity.

Ultimately, the aim of the experiment is to infer **s**_*h*_, which is confounded by the previous biases. Intuitively, the NMF step is designed to remove unspecific reads **x** = **a** *·* **m** *·* **n** from 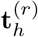, while the HSR step aims to remove the effect of the confounding biases from **y**_*h*_ = **a** · **m** · **s**_*h*_.

The output of DecoDen offers then an approximation of the true hidden signals of histone modification enrichments **s**_*h*_. Further details on the NMF and HSR implementation in DecoDen are provided below.

### Preprocessing

DecoDen has been implemented in Python. For preprocessing, it requires a CSV file specifying details about the sequencing data, such as BAM file location, experimental condition and whether it is a treatment or control file. For each treatment condition, a single fragment length is estimated and reads are extended in the 5’ to 3’ direction. For input control data, the fragment length is estimated as the median of treatment fragment lengths and reads are extended equally in both directions. Next, each BAM file is normalized to have a total library size of 25 million reads. Average coverage is computed for contiguous non-overlapping genomic bins of the specified size (default size is 200). For analyses with real experimental data, the *Low Mappability* regions from the ENCODE blacklist [20] are removed. Deeptools [21] and Samtools are used internally for the manipulation of BAM files. Preprocessed data are written to disk as numpy [22] matrices, with each condition written to a separate file.

### Non-negative Matrix Factorisation

Non-negative matrix factorisation (NMF) is a general methodology to decompose a given matrix V with dimensions *G* × *N* into two matrices *W* and *K* with dimensions *G* × *l* and *l* × *N* respectively such that:

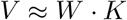

where · denotes the scalar product. The motivation for using NMF is to distinguish histone-specific signals from unspecific additive noise. To achieve this, all ChIP and input measurements are analyzed simultaneously. NMF is used to estimate two quantities - First, consolidated genome-wide signal *W* for each assay *h* and second, a mixing matrix *K*, specifying the composition found in each replicate *r*. Since this step explicitly uses replicates to share information across samples, experimental noise among samples is minimized.

In DecoDen, the matrix *V* ∈ ℝ^*G*×*N*^ consists of a concatenation of treatment and input samples,

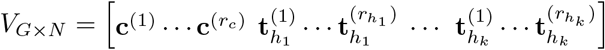

where 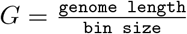, *N* is the total number of samples, *r*_*c*_ is the number of control replicates and each histone modification *h*_*i*_ has a total of 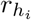 replicates.

The *l* parameter is chosen as the sum of total experimental conditions present in the dataset. For example, in the case where there are ChIP-Seq experiments for two histone modifications with a control, *l* = 3. The learned signal matrix

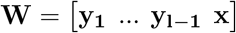

thus contains specific signals for each histone modification **y**_*h*_ = **a** *·* **m** *·* **s**_*h*_, and an additional chromatin bias signal **x** = **a** *·* **m** *·* **n**, containing valuable information about common biases **a** *·* **m**. The mixing matrix *K* contains coefficients, such that the the original measurements 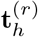 can be reconstructed as linear combinations from **y**_*h*_ and x. The biases captured in the **x** and **y**_*h*_ signals affect both control and ChIP samples, and **x** is thus used in subsequent steps to de-bias the assay-specific signals.

Non-negative matrix factorization is NP-hard in general, and convergence heuristics are dependent on initialization [23]. As the computational cost of NMF is heavily influenced by the sizes the matrix to factorize [24], a heuristic procedure was devised to perform the factorization efficiently and with interpretable structures. 100, 000 genomic bins are sampled from the data matrix as a training set. This training set is used to calculate a mixing matrix with a specified structure, which is then fixed and used to extract the signal matrix from the genome-wide data.

To ensure that the deconvolution step captures the desired information, the matrix factorization operations used to calculate the mixing matrix are split into three steps. Firstly, only the signals for the control replicates are factorized, which results in learning a cell type-specific background and the mixing matrix parameters for the input control samples. Subsequently, this background signal **x** is fixed and the mixing coefficients 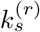 are extracted for the treatment sample that minimize the loss function, to “explain” as much of the treatment signal as possible using only the control signal. Afterwards, the treatment signal is subtracted from this background component and the remaining mixing coefficients jointly with the histone-specific signals **y**_*h*_ is estimated, to explain the remaining information. And finally, the hard constraints on the mixing matrix structure are relaxed to share information across experimental conditions, and to capture the imperfect specificity of antibodies. In this step, we perform NMF optimization using the previously extracted matrices as initialization values; the optimization alternates between fixing the mixing matrix to optimize the signal matrix, and vice versa. This alternated optimization has shown empirically to better preserve the inductive biases encoded in the structure of the factorization procedure. The optimization of all the factorization steps is carried out by minimizing the KL Divergence as loss function.

In summary, the learning scheme is set up in such a way as to explain most of the genomic bins in the ChIP samples with the input sample. This is motivated by the fact that enriched regions constitute about 5-10% of the genome and most of the reads constitute the background signal. Hence, the cell-type specific chromatin background signal *x* in the *W* matrix of NMF contains information common to all replicates and conditions.

As an additional post-processing step, the signals in *W* are normalized so that the 98th percentile is equal to 1, and are multiplied by the corresponding components of *H* to maintain the reconstruction of the original replicates. This additional design choice is intended to concentrate the replicate-specific scaling factors 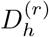 (e.g. due to varying library sizes) in the mixing matrix coefficients, and to make the signals comparable. The choice of percentile to align to a unit value is made to avoid excessive influence of outlier values.

### Half-Sibling Regression

Half-sibling regression [19] is a method inspired by causal inference to remove the effect of confounders. For instance, mappability **m** is a sequence-specific bias, which varies over the genome and confounds measurements at different genomic regions. It is a measure of the number of unique genome positions that a *k*-mer can be aligned against. Another known confounder in estimating the true enrichment of a histone modification is chromatin accessibility **a**. For instance, the highest values for ChIP tracks are higher for histone marks associated with open chromatin when compared to those associated with closed chromatin (See Figure S4 in the Supplementary Material). For a given cell type, this confounder is fixed, and affects both the control sample and the histone-specific signals. In our approach, we model a cell-type-specific confounder as a combination of accessibility and mappability (Refer to the directed graph in Figure 1 C).

In addition, we make the simplifying assumption that the true underlying histone modification signal is independent of the measurement of the cell-type specific signal from the NMF step, implying **s**_*h*_ ╨ **x**. This allows us to use half-sibling regression to remove the effect of confounding factors between the ChIP signal and the cell-type specific background signal. The true histone modification signal **s**_*h*_ is estimated as

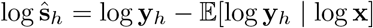

As a design choice, we estimate 𝔼[log **y**_*h*_|log **x**] using a linear regression model.

The application of HSR to logarithm-transformed signals allows us to filter multiplicative noise with better numerical stability. Since subtraction in the log scale translates to division in the linear scale, this is similar to computing fold-change. The estimated signal **ŝ**_*h*_ is recovered by exponentiation of the estimate from half-sibling regression.

Since the HSR step is equivalent to dividing by a confounding factor, certain genomic loci may present extreme values due to a background signal that is essentially 0. To avoid this pitfall, we bounded the divisor to a minimum value of 0.1 (i.e. capping the predictions of the fit in log scale to log(0.1)), essentially ensuring that the signal is at most amplified by a factor of ten.

Non-negative matrix factorization and half-sibling regression are carried out using the Python package *scikit-learn* [25].

### Identifying enriched regions

We use the estimated fold-change from the previous Half-Sibling Regression step and a fixed threshold to identify regions of enrichment. Regions that are close together are merged, and extremely small enriched regions are eliminated. Since no significance tests are done, there is no requirement to correct for multiple testing.

### Training and Validation

In the case of experimental datasets, to evaluate the components of the mixing matrix that describe the proportions of each signal that compose the measured replicates, we select 400,000 genomic bins (bin-width of 200 bp) as training set from regions with average read coverage more than 1. We employed L2 regularization on both the signal matrix and the mixing matrix; to determine the weights of the regularization term in the loss function, we selected 10,000 additional bins to use as validation set to choose a sufficiently high regularization that would not adversely impact the reconstruction of the signal. The final configuration of the algorithm used a weight of 0.01 for the L2 regularization of the signal matrix and 0.001 for the regularization of the mixing matrix. For simulated data, the same regularization parameters were used. Since the simulated genomic region is much smaller than experimental data, the number of genomic bins for training was 5000.

### Data

#### Simulated Data

Since true unbiased histone modification enrichment for experimental data are unknown, synthetic data using the ChIP-Seq simulation tool ChIPsim [26] was generated. ChIPsim is a flexible framework that enables simulation of a wide variety of ChIP-Seq experiments. A custom Markov process was used to assign peak and background regions (Figure S5). Data was simulated for 10 sets of experiments, each with three types of histone modifications (narrow, broad and random peak) and three replicates each. The lengths of background, narrow, broad and random peaks are of 1000 bp, 450 bp, 1200 bp and 750 bp respectively. Details of the simulation procedure are in Section S2 in the Supplementary Material.

#### Experimental Data

##### E114-Jung

Deeply sequenced ChIP-Seq data from Jung et al. [27] measured on the A549 Lung Carcinoma cell line, was used for histone modifications H3K4me3 and H3K27me3 (three technical replicates each) and the whole cell extract (WCE) input sample (two replicates). Raw read data was aligned to the Hg19 reference genome.

##### E114-Roadmap

Raw read data for E114 cell line was obtained from the ENCODE platform and aligned to the Hg19 genome. The two histone modifications H3K4me3 and H3K27me3 had 3 and 2 replicates each, with each treatment sample having a corresponding control sample. Control samples from both assays were used.

##### EN-Tex

DecoDen was run on data for two individuals (37M and 51F) for two tissues (transverse colon and spleen) and for the following histone modifications - H3K4me3, H3K36me3, H3K4me1, H3K27ac and H3K27me3.

Further data processing details (Section S3) and accession numbers (Table S2) are provided in the Supplementary Material.

#### Data Preprocessing

A detailed summary of all datasets used with assays and number of replicates is provided as an additional file. The following procedure was followed when performing alignment of raw sequencing data. SAMtools [28] was used to align the data to the reference genome. Picard was used for quality control of the aligned reads. In particular, reads failing vendor quality checks, those which are PCR duplicates and unmapped reads were removed.

To generate a genome-wide enrichment signal from MACS2, the `bdgcmp` command was used to compare the lambda and treatment tracks obtained from `callpeak` command. This procedure generates a genome-wide −log_10_ p-value enrichment signal in bedgraph format. The Chipseq R package [29] was used to bin this signal for equivalent comparison with DecoDen.

## Results

### DecoDen accurately reconstructs the hidden signal

Since ground truth of histone modification enrichment is unknown for real datasets, DecoDen’s ability to infer true enrichment was first assessed with simulated datasets. Further, DecoDen was quantitatively compared to existing peak calling methods MACS2 [30], GEM [31], MUSIC [32], SICER2 [33], BCP [34] and JAMM [35] (Details in Table S1). Our results show that DecoDen performs competitively in identifying enriched regions in simulated data sets. Most notably, we find that within enriched regions the reconstructed values by DecoDen preserve true coverage patterns (See Figure S6 and Section S2 in Supplementary Material).

Next, we used data from a deeply sequenced experiment from the A549 Lung Carcinoma cell line (equivalent to cell type E114 in the Roadmap Consortium) [27]. This data set will be referred to as E114-Jung in the rest of the manuscript. For this experiment the authors sequenced each sample, including the input background, to generate more than 100M uniquely mapping reads. The authors additionally estimated, that 40M reads would be sufficient to reach saturation. We ran DecoDen run jointly for samples from H3K4me3, H3K27me3 assays and whole cell extract (WCE) as control. Figure 2 demonstrates how the shape of the H3K4me3 enrichment is conserved while eliminating cell-type specific bias at 1.225 Mb. Several other examples highlighting the utility of DecoDen in scenarios where there are a larger number of experimental conditions and replicates available can be seen in Figures S7, S8, S9 and S10 in Supplementary Material.

**Fig. 2:**
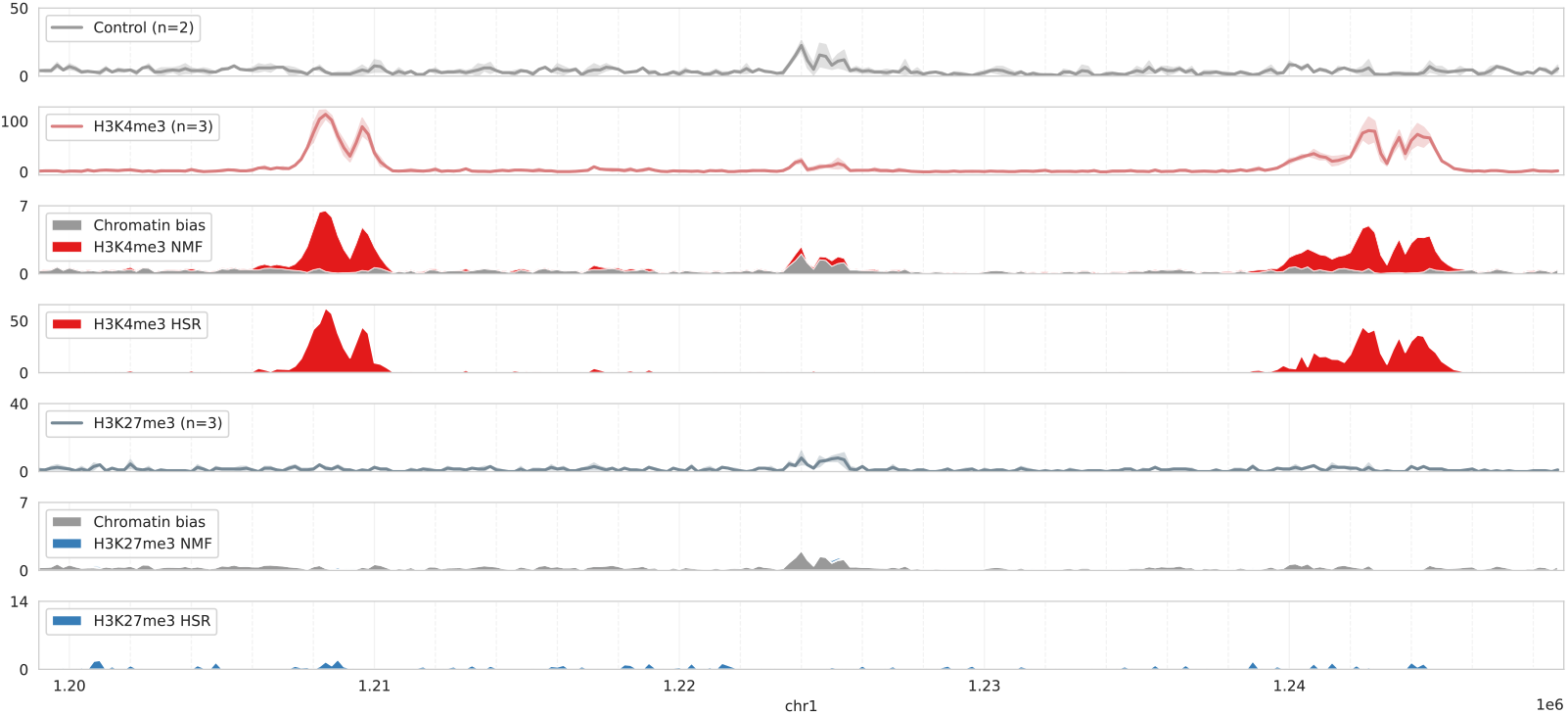
Example DecoDen Reconstruction of H3K4me3 and H3K27me3 enrichment from the E114-Jung Dataset: Measurement data are shown as a solid line with a shaded region around it representing the maximum and minimum read coverage. For each histone modification, the measurement is decomposed into chromatin bias (in grey) and histone modification specific components (in blue and red). While the NMF step reduces noise among replicates, the HSR step eliminates cell-type-specific bias as seen in the region around 1.225 Mb. The shape of the peak is preserved as seen in the H3K4me3 peak on the left side.

The mixing matrix from DecoDen’s deconvolution step can be used as a qualitative diagnostic to assess antibody quality. By analysing publicly available datasets from [36, 37] that use different antibodies to measure the same histone mark, H3K27me3, we reveal differences in antibody specificity directly from the mixing matrix, which align with the specificity determined by the Chromatin Antibodies database [38, 39] (See Figure S11).

### DecoDen minimises the effect of confounders

To determine the true enrichment of a genomic position, it is desirable to remove biases that arise from the experimental process itself, which are ideally captured in the control sample. Figure 3A and B are a compilation of 2D histograms showing the distributions of the average coverage of control and treatment replicates for the histone modifications H3K4me3 and H3K27me3 in the E114-Jung dataset. Notably, the dynamic range for control and signal coverage values are comparable, and the histogram of average coverage present pronounced density across the diagonal: In these regions, count values for the assay are high, but so are control values, indicating a strong correlation between control and signal. In addition, the H3K4me3 data exhibits genomic bins with high relative enrichment along the y-axis, characterized by low control and high assay coverage. For H3K27me3 replicates, relative signal enrichment above background is less pronounced, indicating that the observed sequencing depth may not be sufficient to reliably detect this broad and widespread mark across the genome. This difference arises because the two histone marks have distinct genomic distributions: H3K27me3 spans a large fraction of the genome, whereas H3K4me3 is restricted to a limited number of loci. As a result, the effective local coverage at enriched regions is much lower for H3K27me3 than for H3K4me3. The challenge is further compounded by the fact that H3K27me3 is frequently located in densely packed heterochromatin regions, which are inherently difficult to sequence and yield a lower signal-to-noise ratio. Together, these factors mean that achieving comparable data quality requires a substantially higher sequencing depth for H3K27me3 (and other broad, repressive marks) than for H3K4me3 (See Figure S12). These differences in data quality inevitably affect the performance of downstream computational analyses. Nevertheless, after applying DecoDen, the correlation between input and treatment samples is effectively eliminated. Moreover, DecoDen signals show stronger correlations with assay replicates while minimizing correlation with control samples (Figure 3C).

**Fig. 3:**
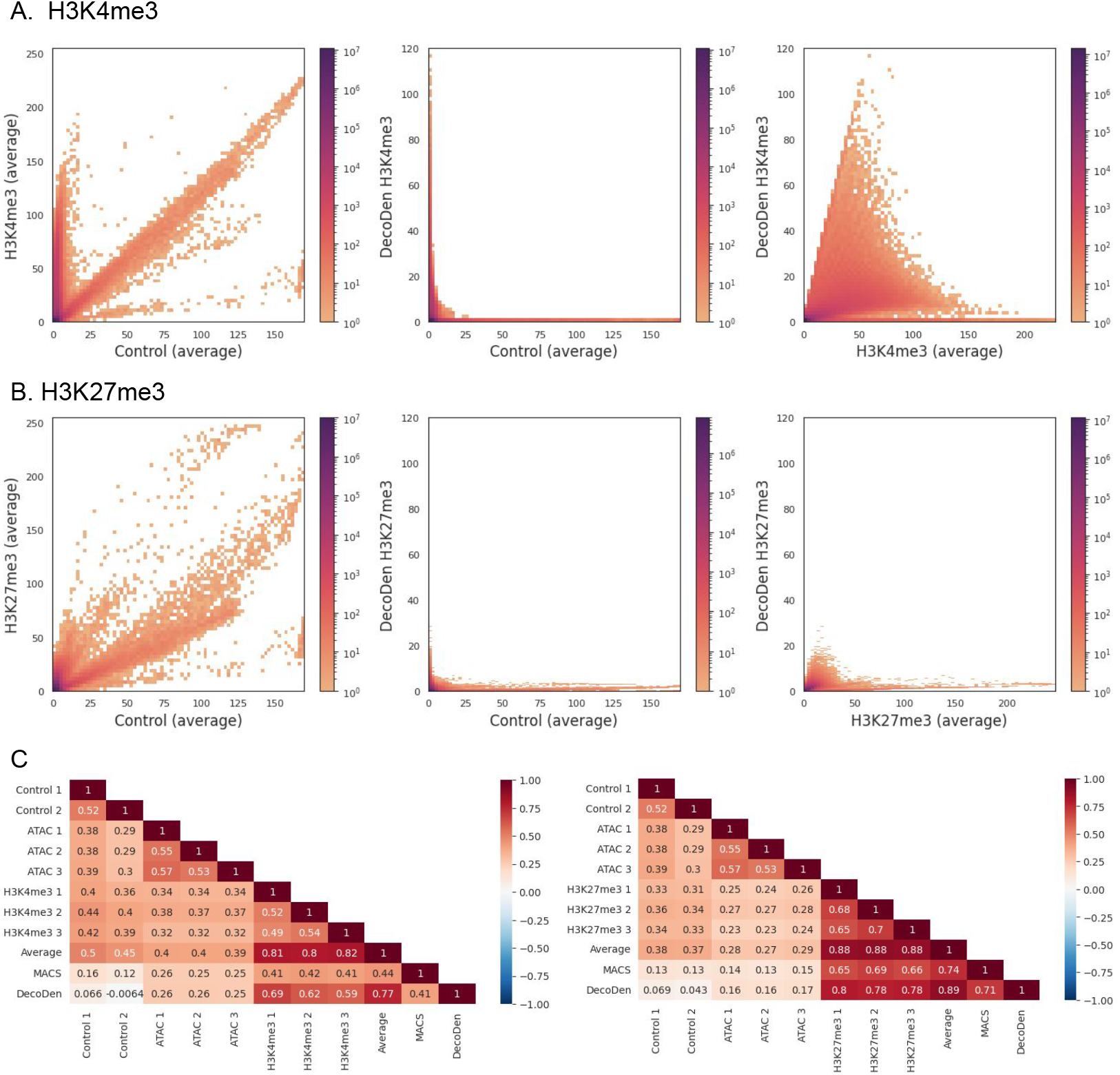
Effect of confounders is minimized: (A) and (B) Histograms comparing the coverage between control and treatment samples are shown for histone marks H3K4me3 and H3K27me27 respectively. As seen in the first columnm, since the local chromatin state influences measurement, treatment and control samples are highly correlated in both histone marks. H3K4me3 has more prominent regions of enrichment than H3K27me3 as seen from the density of genomic bins with high treatment coverage and corresponding low control coverage. DecoDen eliminates this correlation to recover true enrichment as seen in the second and third columns. (C) Heatmaps showing genome-wide correlation demonstrate that while DecoDen signal minimises the correlation with control samples, it enhances the correlation with treatment replicates. Average treatment sample coverage and the p-value signal obtained from MACS are used as a baseline. The E114-Jung dataset has been used here.

When the enrichment signal was independently compared to chromatin accessibility (measured by ATAC-Seq), the correlation was found to be lower than the correlation between assay samples and chromatin accessibility. Specifically, the correlation between H3K4me3 and ATAC-seq (0.32–0.38) was comparable to that between ATAC-seq and the input control (0.29–0.39), and only slightly higher than with H3K27me3 (0.23–0.29). This indicates a shared bias across both ChIP-seq and input samples that is collinear with ATAC-seq, consistent with the fact that open chromatin is inherently easier to sequence than closed chromatin, independent of the targeted histone modification.

Importantly, after applying DecoDen, the correlation between ATAC-seq and the repressive mark H3K27me3 was reduced to 0.16–0.17, while the correlation with the activating mark H3K4me3, which is expected to overlap with open chromatin, remained relatively high at 0.25–0.26. This demonstrates that DecoDen effectively removes sequencing-related biases while preserving true biological enrichment (Figure 3C). DecoDen performs comparably to MACS2 under read subsampling as shown in Figure S13.

### Comparing histone modification measurements across labs

To compare DecoDen’s signal output across experiments, we analysed two independent datasets that measured the same histone marks in the same cell line: the E114-Jung dataset and the corresponding data from the Roadmap Epigenome Project (referred to as the E114-Roadmap dataset). Both datasets include samples for H3K4me3, H3K27me3, and input controls. As seen previously, a strong correlation is present in both raw datasets and their controls indicating that the primary shared signal between experiments is chromatin bias. Consistently, the chromatin bias estimated by DecoDen also correlates strongly across experiments (Figure 4A and B). After correcting for this bias by using either MACS or DecoDen, genome-wide correlations between experiments decrease, suggesting that histone modification patterns are more variable across experimental conditions than initially apparent. This lower post-correction correlation may arise from differences in experimental protocols or antibodies, or it may reflect the inherently dynamic nature of individual histone modifications, which are less stable than overall chromatin architecture and may capture epigenetic responses to distinct external cues during the cellular life cycle. Further, the correlation between DecoDen estimated signal is much higher for H3K27me3, a broad mark, than for H3K4me3, a narrow mark (Figure 4C, D and E).

**Fig. 4:**
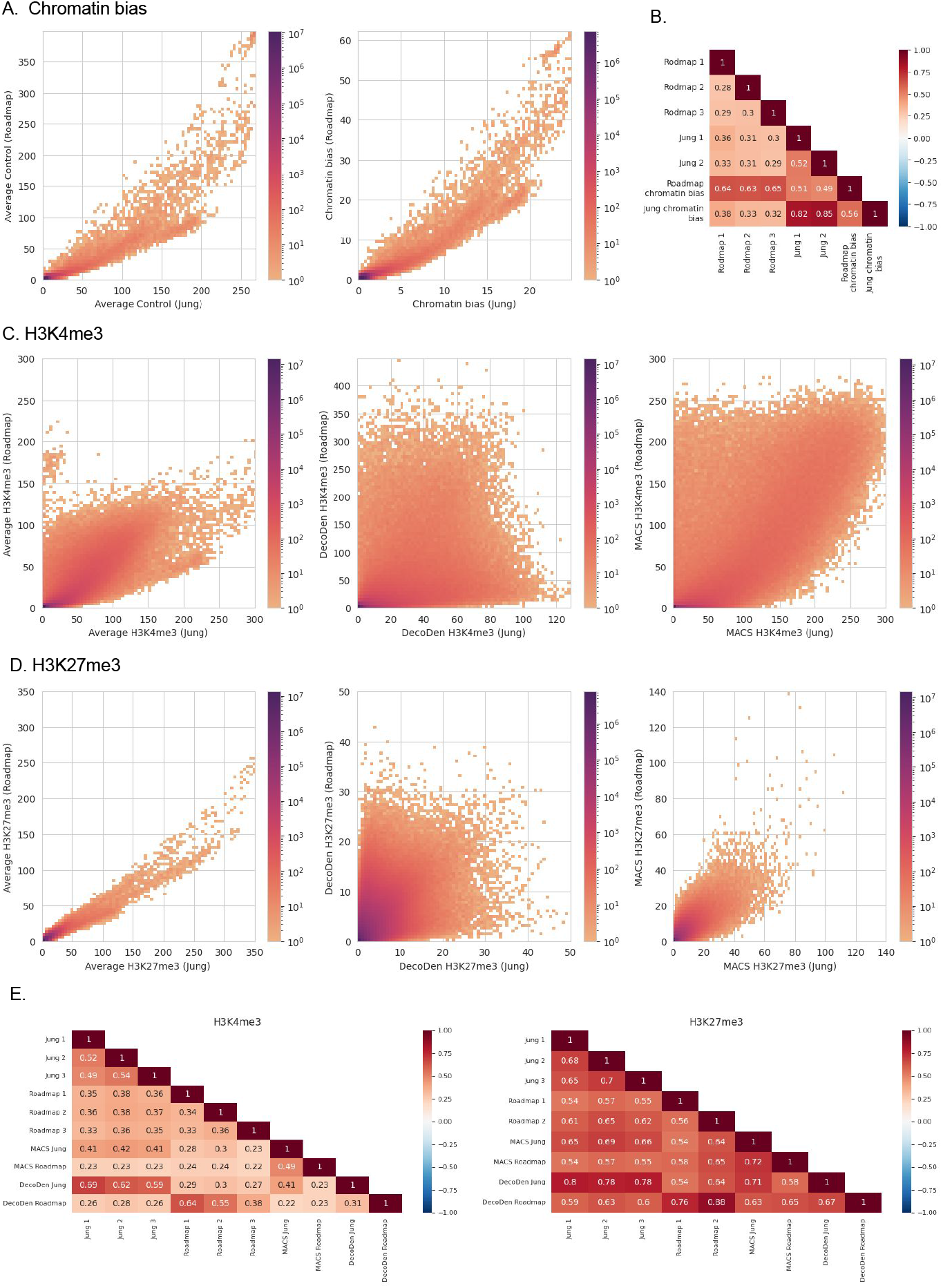
Data comparison between labs: 2D histograms of the average control coverage and DecoDen estimated chromatin bias in (A) and corresponding correlation heatmap in (B) demonstrates the high similarity of bias between both labs. (C) and (D) show 2D histograms for H2K4me3 and H3K27me3 respectively. (E) shows the genome wide correlation between measurement replicates and DecoDen estimated enrichment for both labs.

### DecoDen to study multi-tissue personal epigenomes

Tissue-specific histone modification patterns are influenced by ageing and external factors such as environmental signals. In addition, aberrant histone modification patterns have been associated with abnormal chromatin reprogramming and cancer. Thus, it is important to accurately identify histone modification patterns that are specific to an individual in a given tissue, while reducing biases to better characterize subtle enrichment differences. As a case study, we consider histone modifications in spleen and transverse colon tissue from two individuals of the EN-TEx [18, 40, 41] consortium. Figure 5 shows the 2D histograms for the transverse colon of both individuals and highlights the differences between the observed count distributions between different histone modifications; As in the Jung data set (Figure 3) H3K4me3 and H3K27ac have genomic regions with low control coverage and high treatment coverage along the y-axis. However, in contrast to the earlier examined data sets, the density along the diagonal is much lower, suggesting that the input control sample is severely under-sequenced. The signal enrichment will therefore likely be overestimated, leading to ‘false positive’ enrichment calls. In the case of H3K36me3, H3K27me3 and H3K4me1 signal coverage is itself low, and likely not saturated, suggesting that the sensitivity with which significant enrichment can be detected is expected to be low. Applying DecoDen we can share information across the different assays, thus improving the estimation of the background signal. We are thus able to successfully remove correlation with the estimated chromatin bias from the histone modification tracks (Figure 5 A and B). However, as the control sample and most assay samples appear to be under-sequenced, the resulting output tracks contain estimates of the true histone modification enrichment.

**Fig. 5:**
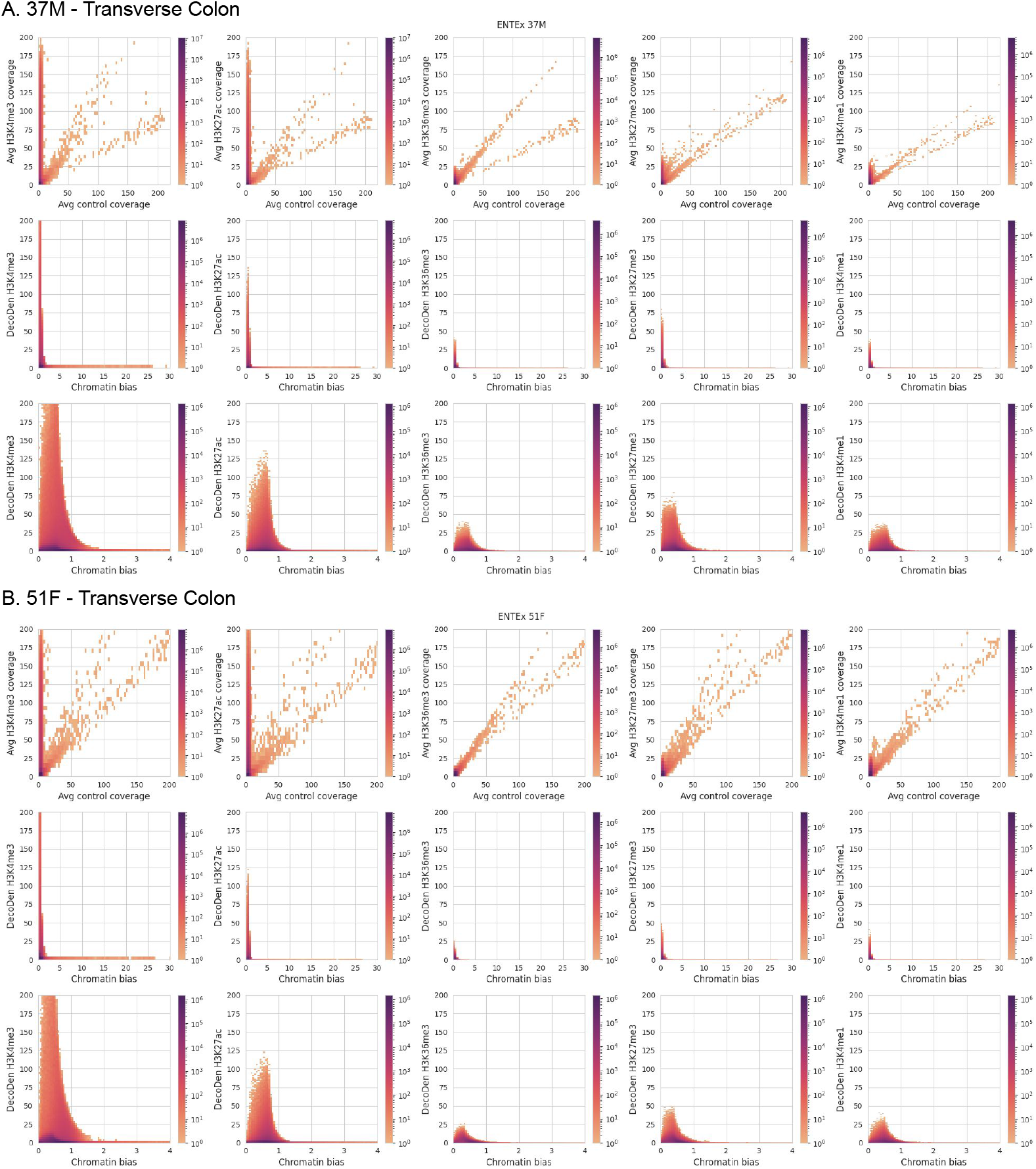
DecoDen removes correlation between control and treatment samples in personal epigenomes: Here, 2D distributions for two individuals from the ENT-Ex dataset are shown. The first row of each sub-figure shows the distributions for binned coverage for input vs assay data. Middle row shows DecoDen estimated histone modification enrichment vs learned chromatin bias. Last row shows enlarged regions of the histograms.

In Figure 6, we compared two individuals based on the DecoDen output and found that the estimated tissue-specific chromatin bias is more strongly conserved between individuals than the assay tracks, despite the low sequencing depth of the control samples. In contrast, greater inter-individual differences were observed for the corrected histone modifications, particularly for H3K4me3 and H3K27ac, the arguably highest-quality tracks. This suggests that individual histone modifications may be less stable and more dynamic, whereas the chromatin landscape, shaped by the interplay of multiple epigenetic factors, remains highly robust within a given tissue context.As before, it is important to note that a higher correlation between different individuals measured using the same assays (e.g., H3K4me1) does not necessarily indicate high reproducibility or data quality. Instead, it primarily reflects the strong and overriding influence of chromatin bias in the assay tracks, as demonstrated by the 2D histograms of the average coverage of measurement data in 6A. This underscores the need for careful correction of such biases when interpreting inter-individual variability in chromatin features. Without proper adjustment, these biases could obscure true biological differences and lead to misleading conclusions about the stability or dynamical nature of these histone modifications versus the broader more robust chromatin landscape.

**Fig. 6:**
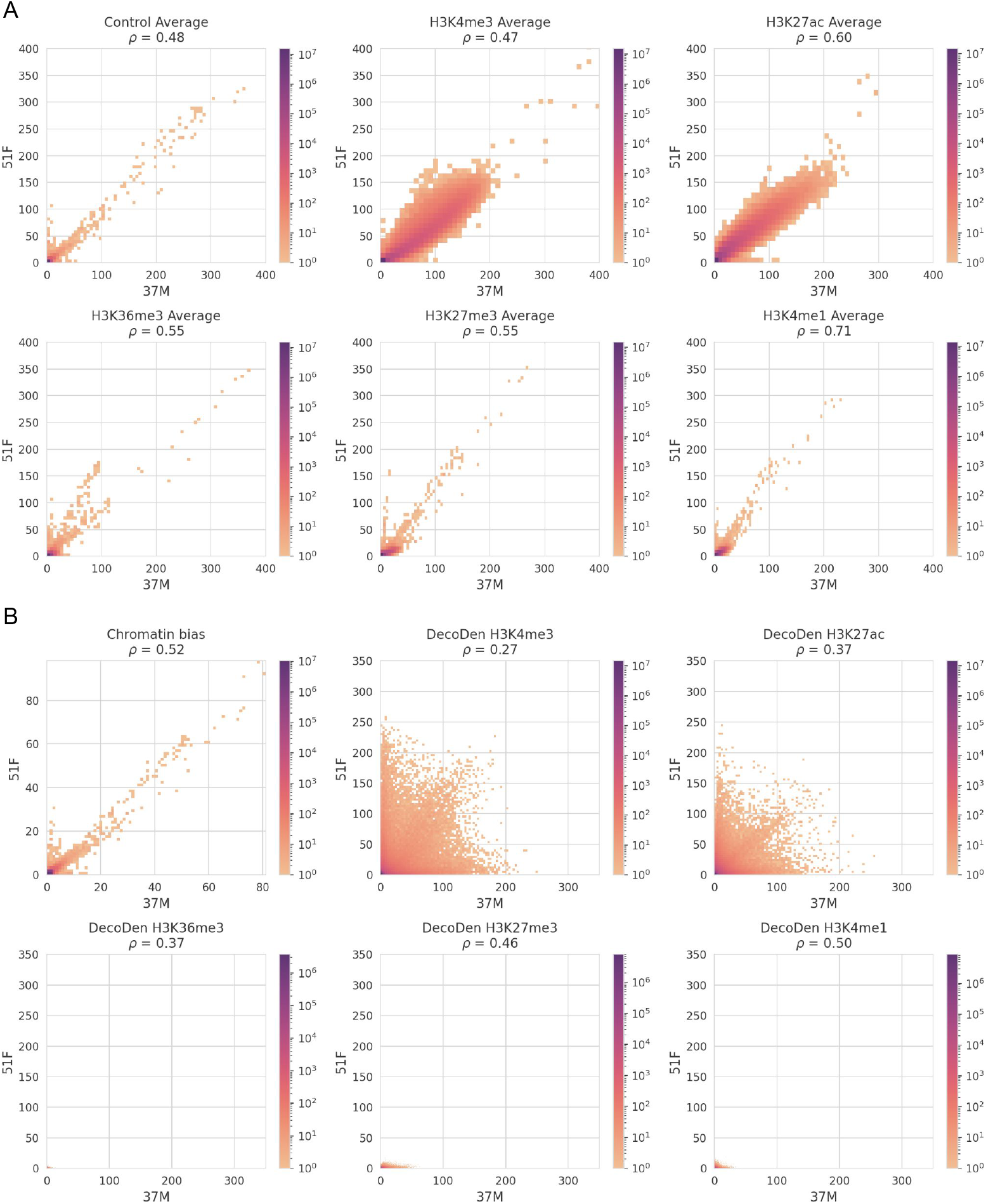
Comparison of DecoDen signal across individuals (transverse colon): (A) shows 2D histograms of average coverage for control and different histone marks across individuals 37M and 51F from EN-TEx dataset, while (B) shows corresponding 2D histograms for DecoDen computed genome-wide signals of histone mark enrichment and chromatin bias. Measurement replicates are correlated across individuals. While chromatin bias is well correlated, histone modifications show a wider range of values. Remarkably, those histone modifications that show a better signal enrichment in average genomic treatment/control coverage are less correlated across individuals than those histone modifications that have a worse genomic treatment/control coverage.

Overall, we observe substantial variability in data quality across histone modifications, likely due to differences in antibody specificity and library sizes, which warrant further investigation. While DecoDen generally enhances enrichment signals, its effectiveness remains constrained by the quality of the input data. Computational methods cannot fully compensate for poor data quality, as seen in histone modifications such as H3K36me3, H3K27me3, and H3K4me1.

Figure 7 examplifies individual-specific H3K4me3 peaks that are more prominent in individual 37M but less pronounced in 51F within the transverse colon. Notably, the replicates for individual 51F exhibit higher variance, leading DecoDen to estimate a lower enrichment value.

**Fig. 7:**
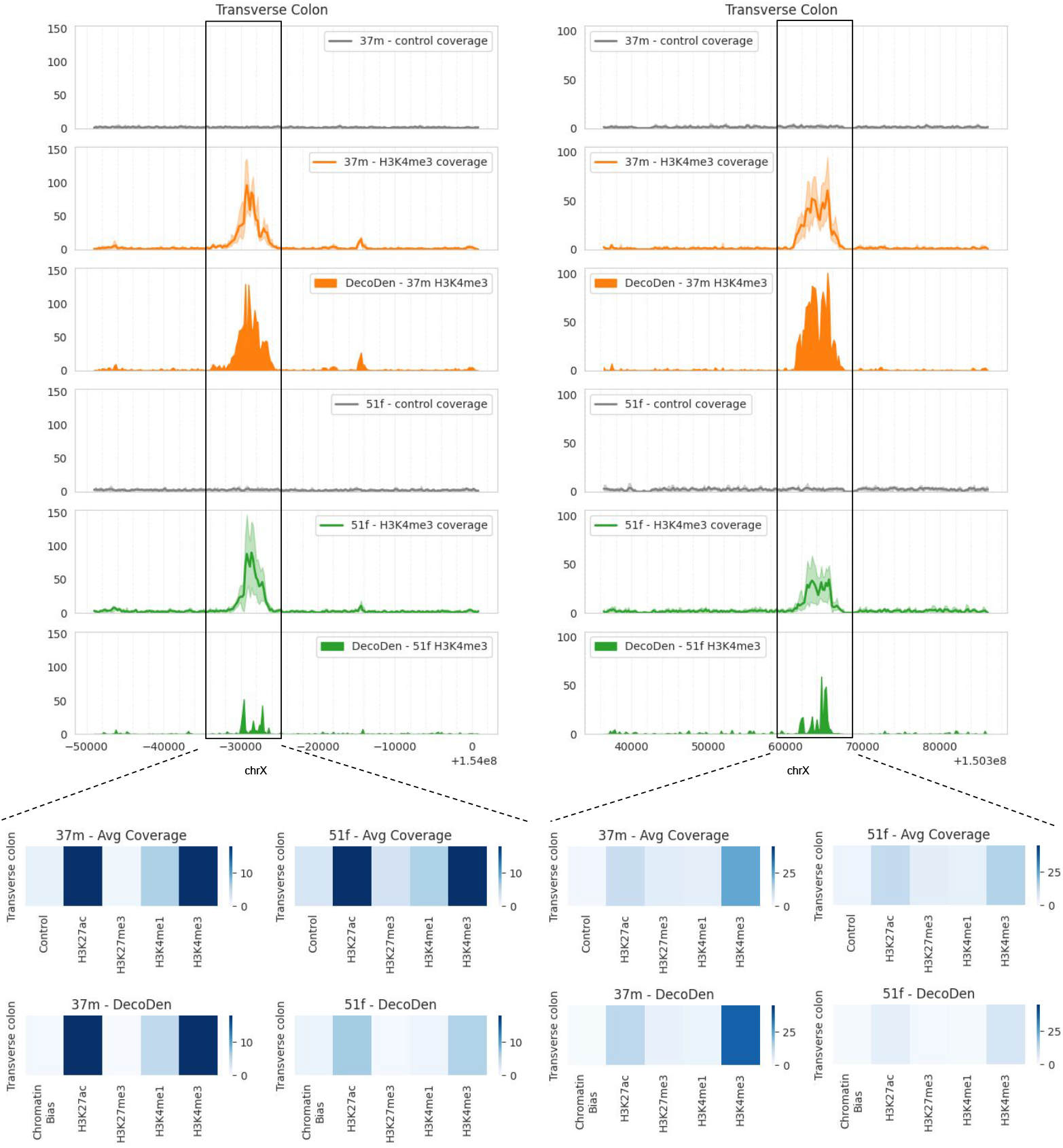
Two examples of genomic loci in transverse colon where a H3K4me3 peak is more prominent in 37M compared to 51F. The histogram below shows the average coverage over different histone modifications in the highlighted genomic region.

## Discussion

DecoDen (Deconvolve and Denoise) leverages replicates and different experimental conditions in histone modification ChIP-Seq experiments to reduce systematic noise and cell-type-specific biases. There are two major steps: non-negative matrix factorization (NMF) to deconvolve the histone-specific signal from the background, and half-sibling regression (HSR) to correct the sources of bias shared between different conditions and replicates.

NMF is used with a custom initialization scheme to optimize the estimation of cell-type-specific signals and histone modification-specific signals. The mixing matrix generated during the NMF step is biologically interpretable, quantifying the contribution of histone-modification-specific and cell-type-specific signals in the ChIP sample. This matrix can also be used to qualitatively evaluate the specificity of antibodies under different experimental conditions. By analysing genome positions collectively rather than independently, the approach operates on a global scale rather than a local one. Additionally, the joint analysis of all samples allows for information sharing across samples and genomic positions, enhancing the robustness of the results. The HSR step incorporates a causal graph of the data-generating process to remove cell-type-specific confounders, such as chromatin accessibility and mappability. This enables the joint analysis of histone modifications that require varying saturation depths.

In ChIP-Seq experiments, treatment and control samples are typically highly correlated. However, the genome-wide enrichment signal derived from DecoDen exhibits a lower correlation with control samples compared to other methods, highlighting its effectiveness in de-biasing ChIP samples. By mitigating confounding effects caused by DNA fragmentation, sonication, and chromatin accessibility, DecoDen recovers previously obscured unbiased enrichment signals. This has also been demonstrated through quantitative comparison of DecoDen with other existing methods using simulated data. For maximum effectiveness, DecoDen benefits from an appropriate experimental design. Ideally, data should include at least two different histone modifications, with replicates for each condition; otherwise, the NMF step becomes redundant. Including both activating and repressive marks is particularly valuable, as greater diversity in the profiled histone modifications ensures that the primary shared component is chromatin bias, which DecoDen is designed to minimize. A broader array of histone marks further enhances performance. In addition, deeply sequenced control samples are highly beneficial, as they provide a direct measure of chromatin bias. Ideally, the control samples should achieve non-zero coverage across most of the genome. Further, the mixing matrix derived from the NMF step can serve as a qualitative diagnostic tool for evaluating sample quality, as high-quality ChIP replicates are characterized by a lower weight for the chromatin bias.

Interestingly, comparisons across experiments from different laboratories reveal higher concordance for histone modifications associated with lower-quality data. This is likely due to the pronounced similarity in chromatin bias, as lower-quality ChIP-Seq samples tend to show stronger correlations with input controls.

A similar trend emerges in the analysis of individual epigenomes within the EN-TEx dataset, where substantial variability in data quality is observed across histone modifications. To accurately quantify individual-specific epigenomic differences, further investigation using higher-quality data is essential. The effectiveness of downstream computational models, including imputation methods, is heavily dependent on data quality; thus, robust quality assessment is critical for developing reliable machine learning models in personalized epigenomics. Further, as large-scale deep learning models trained on ENCODE and EN-TEx data are increasingly used to predict histone mark enrichment profiles, our results highlight the need for caution regarding the quality of the input data used to train these models, and consequently on the reliability and accuracy of their predicted histone mark profiles.

In conclusion, we introduce DecoDen, a computational method designed to share information across measurements, thereby enhancing data quality through computational means. The effectiveness of DecoDen is demonstrated by its ability to improve histone modification enrichment, as evidenced by comparisons across laboratories for the same cell type and its application to individual epigenome data from the ENTEx dataset.

## Supporting information

decoden_supplement

decoden_data

## Acknowledgements

The authors thank the International Max Planck Research School for Intelligent Systems (IMPRS-IS) for supporting Tanmayee Narendra and Crhistian de Jesus Cardona. We would also like to thank Tom Owen-Hughes and Kasper Rasmussen for helpful comments and suggestions.

