## Supplementary material for "Towards Personalized Epigenomics: Learning Shared Chromatin Landscapes and Joint De-Noising of Histone Modification Assays": decoden_supplement

### Supplementary Information

#### S1 Confounding by Mappability and Chromatin Accessibility

The main objective of using a control sample in a ChIP-Seq experiment is to estimate the background distribution of genomic reads. This background distribution is influenced primarily by mappability and chromatin accessibility and the control sample is a way to measure this. Mappability is defined as the inverse of the number of unique places that a k-mer occurs in a genome. It is sequence specific and does not change among histone modifications or across different tissues. In other words, for a fixed organism, mappability is the same across different tissues. It is inherently difficult to make claims about enrichment at regions of low genome mappability and any peaks identified at these regions are dubious. On the other hand, chromatin accessibility is tissue specific.

It is important to sequence the control sample to a sufficient depth to ensure that the mappability and chromatin accessibility is estimated correctly. As an example, consider the four control replicates for E004 (H1 BMP4 Derived Mesendoderm Cultured Cells) from the Roadmap dataset. Since these are technical replicates, biological variation is at a minimum. Approximately 1 billion bp have a coverage of 0 in all

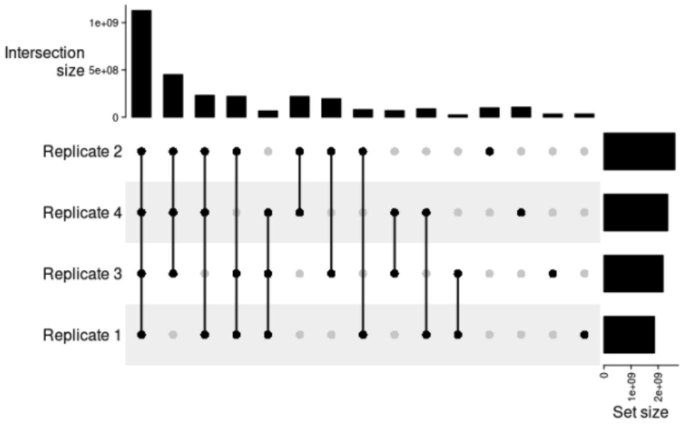


**Fig. S1**: Overlap of regions with zero coverage for E004. E004 has 4 control replicates.

replicates. (See Figure S1 for details), which constitutes more than 16% of the human genome. Hence, at least 16% of the background distribution is systematically underestimated. Similar behaviour is seen among the four technical replicates for E007 (H1 Derived Neuronal Progenitor Cultured Cells), see Figure S2 for details. There is a high overlap among zero coverage regions across tissues, see Figure S3.

Low control coverage also affects activating and repressive marks differently. We plotted the maximum $-\log_{10} p$ for different tracks in the Roadmap dataset

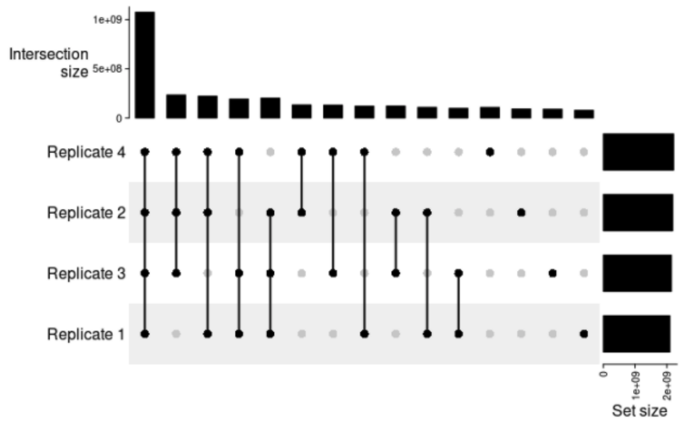


**Fig. S2**: Overlap of regions with zero coverage for E007. E007 has 4 control replicates.

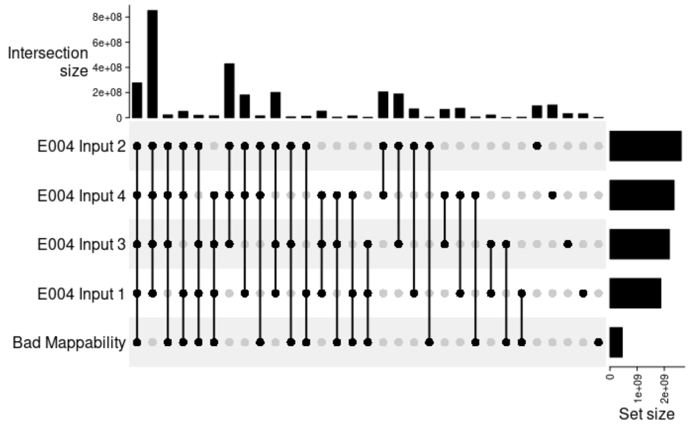


**Fig. S3**: Upset Plot showing overlap of zero coverage regions in control samples among different tissues. Roadmap's consolidated data was used for this comparison. The ENCODE CRG Mappability track for 36-mer was used with a threshold of 0.1 to define regions of bad mappability.

and grouped activating and repressive marks separately. Overall, repressive marks (H3K9me2, H3K9me3 and H3K27me3) have a lower maximum significance compared to activating marks (H3K9ac, H3K4me2, H3K4me3 and H3K36me3) as seen in Figure S4 [42].

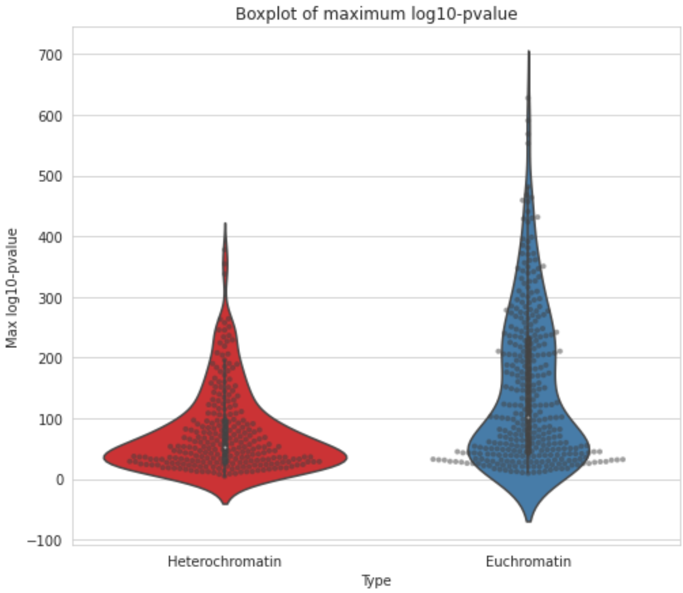


**Fig. S4**: Violinplot of maximum MACS2 $-\log_{10}$(pValue) for histone marks associated with euchromatin versus those associated with heterochromatin. The signal is restricted to chromosome 21 and binned to 25 bp width.

### S2 Simulation Experiments

#### S2.1 Data Simulation

Data for quantitative evaluation is simulated using the *ChIPsim*[26] Bioconductor package. We adapted the procedure described in the package documentation to reproduce the simulations used in [43]. Data was simulated for 10 sets of experiments, each with three histone modifications and three replicates per modification. Details of the simulation are described below.

A genome of length 1 million base pairs (bp) is used for simulation. A Markov process (See Figure S5) is used to assign contiguous genomic regions into four categories - background, narrow peak, broad peak or random peak. The lengths of the background, narrow, broad and random peaks are of 1000 bp, 450 bp, 1200 bp and 750 bp respectively. The process is initialised to a background region and every background $\rightarrow$ peak transition is assigned a probability of 0.05. The transition probability from peak $\rightarrow$ background is 1, ensuring that each peak region is flanked by background regions. Each run of the Markov process assigns different background/peak regions, hence producing a new dataset for evaluation.

Similar to [43, 44], the value for the binding density for background regions is sampled from a Gamma distribution with shape parameter 1 and scale parameter 20. This probability is a measure of how susceptible this genomic region is to non-specific binding events. The corresponding binding density for peak regions is sampled from

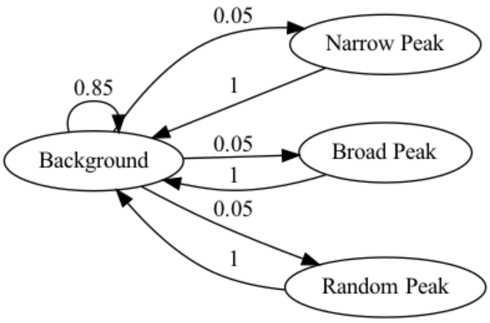


**Fig. S5**: A custom Markov process is used to assign peak/background features to genomic regions. The numbers on edges represent the transition probabilities of the connecting states.

a Pareto distribution with shape parameter 2 and a minimum value of $20t$. The $t$ parameter determines the noise level. This is chosen to be 15, 10 and 12 for Narrow, Broad and Random peak respectively.

The next task is to determine the shape of the binding density. For peak regions, binding densities represent the propensity of the region to bind a protein of interest. For background regions, it is an indication of non-specific binding. Each genomic region is assigned a binding density with the shape of a Beta distribution. The shape parameters $\alpha$ and $\beta$ are sampled uniformly from the interval $[2, k]$ where $k$ is 3, 10, 7 and 8 for background, narrow peak, broad peak and random peak respectively. To simulate a particular experimental condition (input, narrow peak, broad peak or random peak), features that are irrelevant to the experimental condition are replaced with an identical background feature. For example, consider the case where we want to simulate replicates for the narrow peak regions. To do this, all genomic regions of type random and broad peak are replaced by background features with a binding density as the average of the two flanking background regions.

Fragment lengths are assumed to follow a truncated normal distribution with mean length of 200 bp, and minimum and maximum of 150 bp and 250 bp respectively. The probability that a given base pair generates a read is generated from the convolution of the fragment length distribution with the binding density. This resulting distribution is used to sample 36 bp reads across the genome. Reads are written into BED files using *GenomicRanges*[45].

#### S2.2 Running peak calling tools

We first assessed DecoDen's ability to distinguish between true enriched regions and background regions and compared it to the performance of peak calling tools. The six tools used for this comparison are: MACS2 [30], GEM [31], MUSIC [32], SICER2 [33], BCP [34] and JAMM [35]. Further details are provided in Table S1.

While several tools allow to specify different replicates in a single run, only JAMM and DecoDen use replicates separately. MACS2 internally merges all treatment and control replicates respectively. All the peak calling methods were run with different

| Tool | Version | Input | Significance | URL |
|---|---|---|---|---|
| MACS2 | 2.2.7.1 | BED/BAM | Q-value | github.com/macs3-project/MACS |
| MUSIC | 1.0.2 | BED | Q-value | github.com/gersteinlab/MUSIC |
| GEM | 3.4 | BED/SAM | Q-value / FC | groups.csail.mit.edu/cgs/gem/ |
| BCP | 1.1 | BED | P-value | rulai.cshl.ed/BCP/ |
| SICER2 | 1.0.2 | BED/BAM | FDR | github.com/zanglab/SICER2 |
| JAMM | 1.0.8 | BED | Q-value | github.com/mahmoudibrahim/JAMM |

**Table S1**: Table showing the peak calling methods used for the comparison. The official URL for BCP is not reachable. We used a repository copy available in: https://github.com/will-NYGC/bcp. (FC: fold change, FDR: false discovery rate)

values of significance threshold (Q-value, P-value, Fold change, False discovery rate) that allows variation in the number of identified peaks. For each tool, the default value was used for all other parameters. An automated pipeline was written using the SnakeMake library [46] and executed using Conda in an HPC with subq as the job scheduler. See Table S1 for a summary of the different tools used.

Minor modifications were made to JAMM and SICER2 to run them on the simulated datasets. Since GEM only reports exact genomics locations of binding, we used the same approach of [44] where a 200 bp window around these identified binding locations was used to define peaks for comparison with other methods. All code for evaluation, including modified versions of libraries, is in the DecoDen repository at `https://github.com/ntanmayee/DecoDen`.

#### S2.3 Performance evaluation

Each tool was evaluated on 10 simulated datasets, with each dataset containing three conditions (narrow, broad and random) with 3 replicates for each condition. Each set of peaks was compared to the ground truth of binding features using the *findOverlapsOfPeaks* function in the ChIPpeakAnno[47] and Bioconductor[48] package in R[49]. We also disambiguate the metrics by avoiding one-to-many and many-to-one matches between true and predicted peaks, normalizing the results such that only we could see one-to-one matches.

Figure S6 A shows the results of the comparison of different methods. The metrics used are sensitivity, precision and F1 score. Similar to [44], sensitivity is defined as the fraction of the true peaks that overlaps with the significant peaks, precision is defined as the fraction of the significant peaks that overlap with the true binding features and F1 score is the harmonic mean between sensitivity and precision. In most cases, DecoDen performs best among all methods, closely followed by MACS2.

In addition to identifying enriched regions it is important to correctly preserve the amplitude of the enrichment. Figure S6 B-D shows specific examples where there is a mismatch between DecoDen and MACS2. MACS2 is chosen for comparison over other methods as its performance is superior. However, MACS2 tends to distort the shape of the peak while DecoDen faithfully reproduces the enrichment in the final signal. Since MACS2 merges control replicates and uses this to compute the background, the final enrichment signal is strongly influenced by the control replicates.

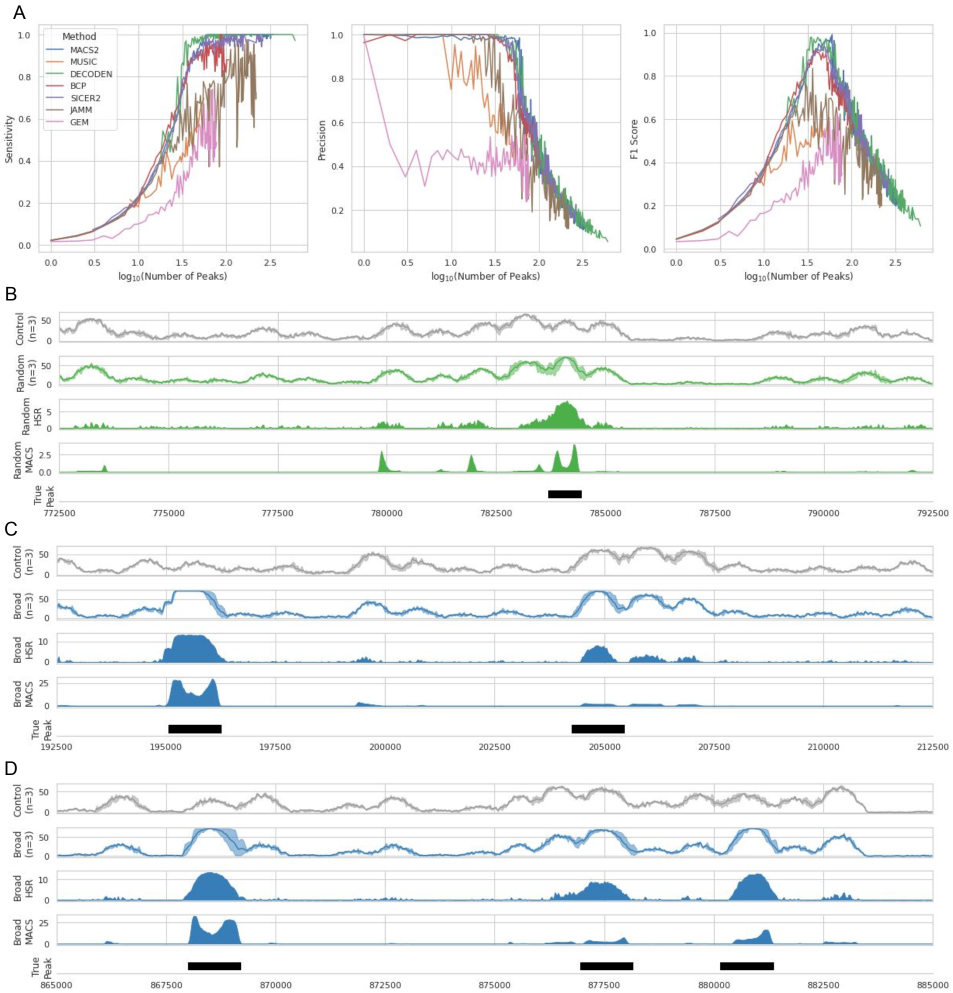


**Fig. S6**: **Quantitative evaluation on simulated data:** A. Sensitivity, precision and F1-score for all methods. Only the mean values are shown for clarity. B-D: Examples of mismatch between DecoDen and MACS2. In each plot, the first two lines show the control and treatment coverage respectively. The shaded area indicates the range of values among replicates. The next two rows show the enrichment signals from DecoDen (experimental condition-specific signal) and MACS2 respectively. The last row shows the true peak region used for simulation. B and C show examples where MACS2 distorts the shape, and D is an example where peaks are missed.

### S3 Experimental Data

Data from the following sources have been used :

- Jung et al: Figures 2, 3, 4, **??**

- Roadmap: Figures 4, S1, S2, S3, S7, ??
- EnTEx :Figures 5, 6, 7, S8, S9, S10
- Antibody specificity data: Figure S11

Further details about experimental conditions and number of replicates have been provided in an additional file.

#### S3.1 Processing for Roadmap Consortium Data

For H1 BMP4 Derived Trophoblast Cultured Cells (E005) from the Roadmap Consortium, the following histone modifications were used - H3K9me3, H3K27me3, H3K4me3, H3K4me1, H3K36me3, H3K27ac and H3K9ac in addition to WCE input. Since DecoDen leverages multiple replicates, the unconsolidated data was used. As E005 is a cell line, the data consisted exclusively of technical replicates. The data set is provided as `tagAlign` files which have been aligned to the hg19 genome.

Since data for E114 is additionally available on ENCODE, the reads were directly obtained and processed according to the pipeline described in Section 2.

#### S3.2 Accession codes

The SRA accession code for deeply sequenced Human A549 cell line is SRP038005.

The accession numbers for the datasets downloaded from the ENCODE portal are in Table S2.

| 51 years, Female, Spleen (Biosample accession: ENCBS363GBQ,ENCBS892ESJ) | | |
|---|---|---|
| **Accession** | **Assay title** | **Target of assay** |
| ENCSR826MTK | Histone ChIP-seq | H3K27me3 |
| ENCSR589DBF | Histone ChIP-seq | H3K4me3 |
| ENCSR831EDZ | Histone ChIP-seq | H3K4me1 |
| ENCSR272CIV | Histone ChIP-seq | H3K9me3 |
| ENCSR711URW | Control ChIP-seq | - |
| ENCSR668GBL | Histone ChIP-seq | H3K27ac |
| 51 years, Female, Transverse colon (Biosample accession: ENCBS555CAN,ENCBS515RIZ) | | |
| ENCSR500QVK | Histone ChIP-seq | H3K4me1 |
| ENCSR384KIB | Control ChIP-seq | - |
| ENCSR488YXS | Histone ChIP-seq | H3K36me3 |
| ENCSR315EZG | Histone ChIP-seq | H3K4me3 |
| ENCSR792VLP | Histone ChIP-seq | H3K27ac |
| ENCSR604QMH | Histone ChIP-seq | H3K27me3 |
| 37 years, Male, Spleen (Biosample accession: ENCBS486GAC,ENCBS511RRL) | | |
| ENCSR091IRL | Histone ChIP-seq | H3K9me3 |
| ENCSR506DCR | Histone ChIP-seq | H3K27me3 |
| ENCSR889RIF | Control ChIP-seq | - |
| ENCSR426VHO | Histone ChIP-seq | H3K27ac |
| ENCSR407UEE | Histone ChIP-seq | H3K36me3 |
| ENCSR556QXD | Histone ChIP-seq | H3K4me3 |
| ENCSR928BGP | Histone ChIP-seq | H3K4me1 |
| 37 years, Male, Transverse colon (Biosample accession: ENCBS990XOM,ENCBS816BKF) | | |
| ENCSR083DXD | Histone ChIP-seq | H3K9me3 |
| ENCSR813ZEY | Histone ChIP-seq | H3K4me3 |
| ENCSR749ABB | Histone ChIP-seq | H3K36me3 |
| ENCSR516QFO | Histone ChIP-seq | H3K4me1 |
| ENCSR643KID | Histone ChIP-seq | H3K27me3 |
| ENCSR479HSY | Control ChIP-seq | - |
| ENCSR640XRV | Histone ChIP-seq | H3K27ac |

**Table S2**: ENCODE Accession List

### S4 DecoDen Examples

### S5 Antibody specificity

Determining antibody specificity aprioiri is a difficult problem. With DecoDen, it is possible to qualitatively assess antibody quality with the mixing matrix of the NMF step. To illustrate this, we chose two antibodies with different affinities for unspecific reads for the histone modification H3K27me3. Both these antibodies have been independently assessed in the Chromatin Antibody database [38, 39]. The two antibodies chosen were Abcam `ab192985` from [36] (7.83% unspecific) and Abcam `ab6002` (59.41% unspecific) from [37]. From the mixing matrices in Figure S11, it is clear that `ab192985` has lower unspecific reads as compared to `ab6002`.

### S6 Narrow versus broad histone marks

The goal of the simulation was to demonstrate the differences in the genomic distributions of H3K4me3 and H3K27me3. Code for the simulation can be found at

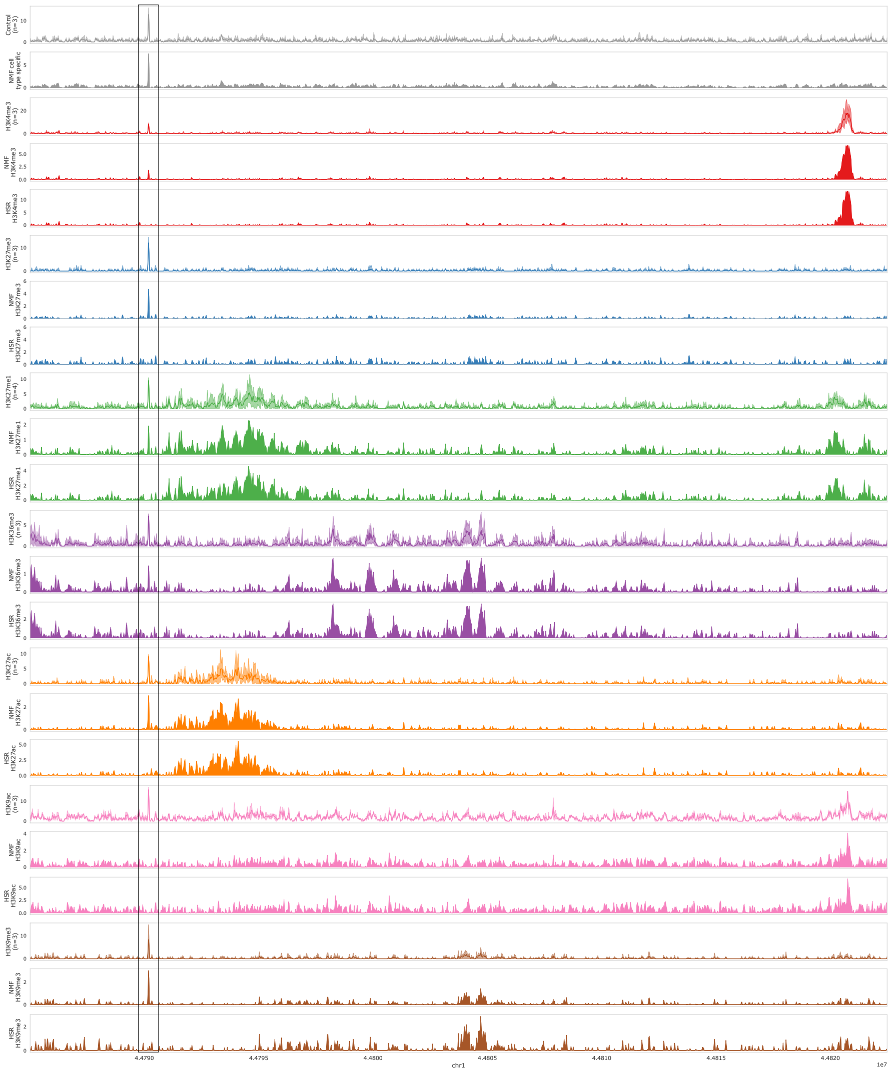

**Fig. S7**: **Reconstruction using DecoDen for H1 BMP4 Derived Trophoblast Cultured Cells (E005) from the Roadmap Consortium:** The first row in each colour shows coverage of replicates after the pre-processing step. The second and third row show the signal after the NMF and HSR steps, respectively. The highlighted spike is identified as a cell-type specific signal in the NMF step and is removed from all histone marks after HSR.

https://gist.github.com/ntanmayee/5b502aa55e0f1d269758e1ffb8fa03f0. The general approach is described below:

1. First a genome length of 1000 bp is defined, with the first 600 bp is heterochromatin, and the rest is euchromatin. Heterochromatin has a lower probability of sequencing.

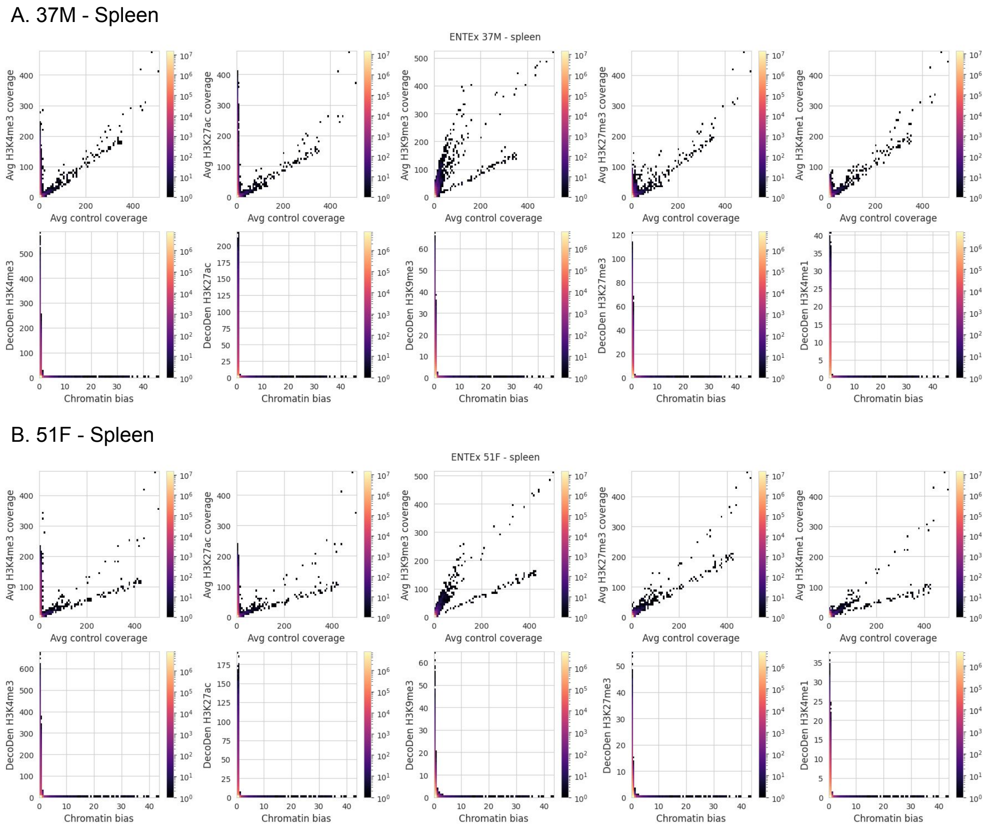


**Fig. S8**: **ENTE-x Spleen for individuals 37M and 51F:** 2D histograms of average coverage of treatment versus control replicates and DecoDen estimated chromatin bias and enrichment.

2. A narrow peak and a broad peak are defined in euchromatin and heterochromatin respectively. The shape of each peak is defined by a beta distribution.
3. Reads are generated for each experimental condition, i.e., control, H3K4me3 and H3K27me3, for a fixed library size. Read coverage is subsequently computed.

The corresponding coverage can be seen in Figure S12, where the broad mark H3K27me3 shows lower coverage than the narrow mark H3K4me3.

### S7 Subsampling of reads

To compare DecoDen and MACS2's performance when reads are subsampled, all replicates from E114-Jung dataset were subsampled at fractions $\{0.1, 0.2, ..., 0.9\}$. DecoDen's estimated enrichment was compared with the binned MACS2's p-value signal obtained from the `bdgcmp` command. The genome-wide correlation at each fraction

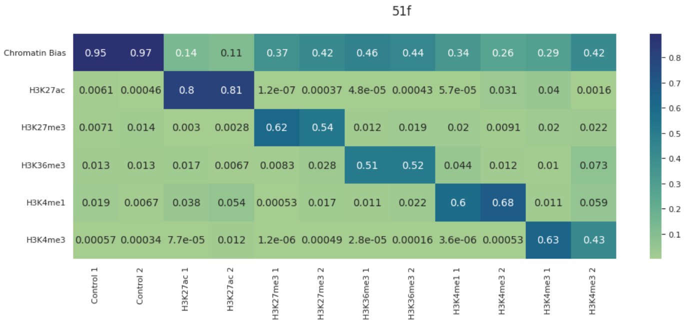


**Fig. S9**: Mixing matrix for transverse colon of individual 37M. Each column is normalised to sum to 1.

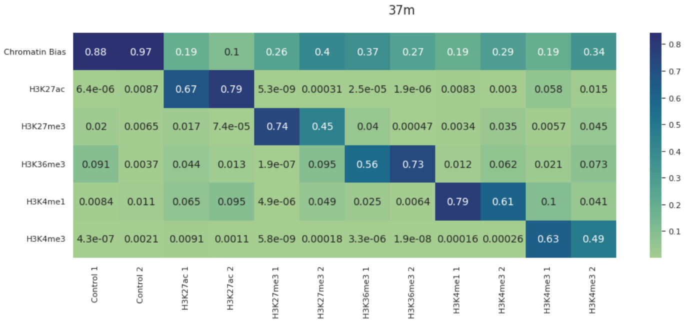


**Fig. S10**: Mixing matrix for transverse colon of individual 51F. Each column is normalised to sum to 1.

was compared to the correlation at 100% of reads. DecoDen's performance is comparable to MACS2, with MACS2 shows a slight improvement over DecoDen's correlation (Figure S13).

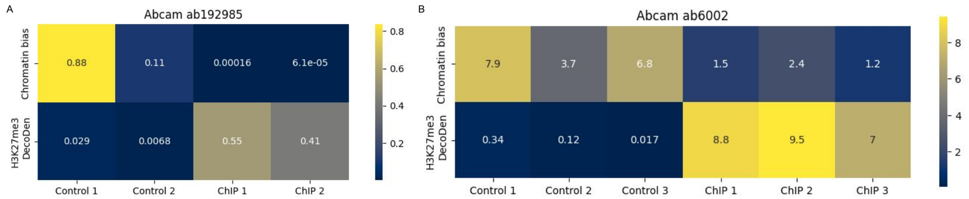


**Fig. S11**: **DecoDen recovers antibody specificity**: The mixing matrices for two different ChIP-Seq experiments measuring H3K27me3 using different antibodies (Abcam ab192985 and ab6002) are shown. As seen from the ChIP columns, antibody ab192985 has a lower percentage of unspecific reads compared to antibody ab6002.

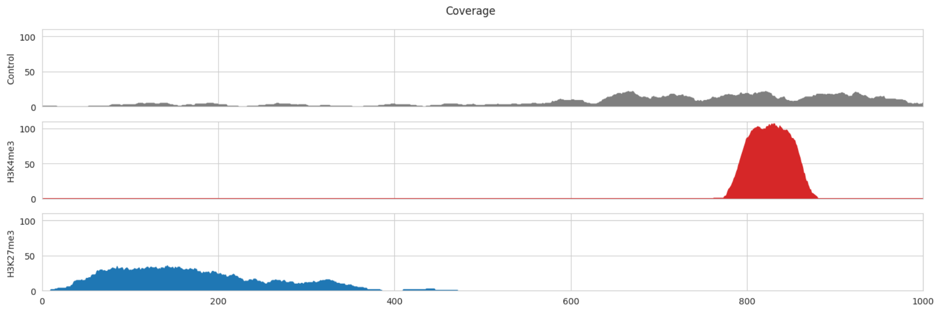


**Fig. S12**: To demonstrate different genomic distributions for H3K4me3 and H3K27me3, reads were simulated. The figure shows a comparison of coverage for input control (top row), H3K4me3 in the second row(narrow, activating mark) and H3K27me3 in the last row (broad, repressive mark).

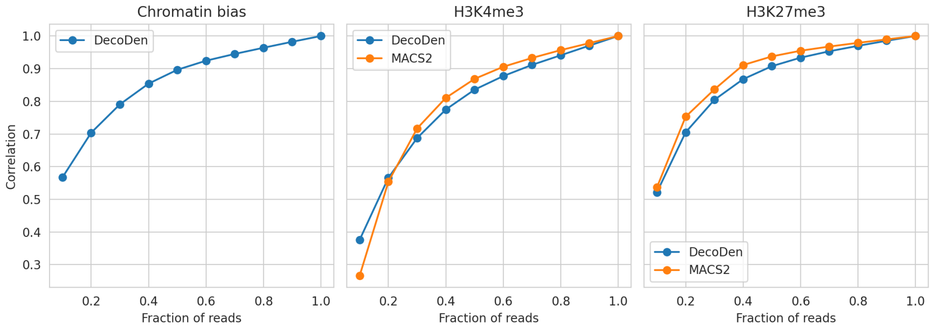


**Fig. S13**: Comparison of correlation of subsampled reads between MACS2 and DecoDen
